# GENA-Web - GENomic Annotations Web Inference using DNA language models

**DOI:** 10.1101/2024.04.26.591391

**Authors:** Alexey Shmelev, Maxim Petrov, Dmitry Penzar, Nikolay Akhmetyanov, Maksim Tavritskiy, Stepan Mamontov, Yuri Kuratov, Mikhail Burtsev, Olga Kardymon, Veniamin Fishman

**Affiliations:** Artificial Intelligence Research Institute, AIRI, Moscow, Russia; HSE University, Moscow, Russia; London Institute for Mathematical Sciences, London, UK; Institute of Cytology and Genetics, Novosibirsk, Russia

## Abstract

The advent of advanced sequencing technologies has significantly reduced the cost and increased the feasibility of assembling high-quality genomes. Yet, the annotation of genomic elements remains a complex challenge. Even for species with comprehensively annotated reference genomes, the functional assessment of individual genetic variants is not straightforward. In response to these challenges, recent breakthroughs in machine learning have led to the development of DNA language models. These transformer-based architectures are designed to tackle a wide array of genomic tasks with enhanced efficiency and accuracy. In this context, we introduce GENA-Web, a web-based platform that consolidates a suite of genome annotation tools powered by DNA language models. The version of GENA-Web presented here encompasses a diverse set of models trained on human data, including the prediction of promoter activity, annotation of splice sites, determination of various chromatin features, and a model for scoring of enhancer activity in Drosophila. GENA-Web is accessible online at https://dnalm.airi.net/

## INTRODUCTION

Understanding the sequence determinants that underpin the functionality of genomic elements is crucial for annotating genomes and interpreting genetic variants. While biochemical approaches to genome analysis can yield direct information about genome functions, these approaches are often constrained by the complexity of molecular systems interacting with nucleic acids. Recently, machine learning-based approaches have emerged as powerful alternatives, capable of inferring a wide range of epigenetic and genomic features from DNA sequences with remarkable accuracy (1, 2, 3). Notably, transformer-based neural networks have emerged as the leading architecture, delivering unparalleled results in the field, as evidenced by various studies and implementations (1, 2, 4, 5, 6, 7, 8).

The utility of machine learning in genomics is significantly enhanced by the application of transfer learning, a method that leverages knowledge gained from one task to improve performance on another. In the genomic context, this is typically achieved by initially pre-training a model on a specific biological task, enabling it to grasp the fundamental patterns and structures within DNA sequences. After this foundational learning phase, the model can be fine-tuned to address a variety of specialized downstream tasks with greater efficiency (4, 5, 6, 7, 8). Although the pre-training phase may demand substantial computational resources, the subsequent fine-tuning process is generally quicker, offering a cost-efficient approach to applying advanced machine-learning techniques in genomic research.

The development of pre-trained DNA language models has seen significant advancements recently, beginning with the pioneering DNABERT transformer (4). Pretrained on the human genome, DNABERT demonstrated its ability to accurately predict promoter activity, splice site localization, and transcription factor binding sites after undergoing task-specific fine-tuning. The promising results achieved by DNABERT have spurred the rapid development of transformer-based pre-trained models, leading to the introduction of BigBird (9), NucleotideTransformer (6), DNABERT-2 (8), and GENA DNA language models (GENA-LMs) (7). These newer models have expanded upon the capabilities of DNABERT, featuring an increased number of parameters and, crucially, the capacity to process longer input sequences. This enhancement in handling extended sequences has proven to be vital for addressing a wide range of downstream genomic tasks (7). Notably, among the publicly available transformer-based pre-trained models, GENA-LMs boast the longest sequence input capability, accommodating up to 32 kb (7). This extended input range enables GENA-LMs to surpass DNABERT in nearly all biological tasks, showcasing the significant impact of input sequence length on the performance of DNA language models in genomic research.

While fine-tuning DNA language models has proven effective for addressing a variety of biological tasks, there’s a notable gap in the accessibility of these fine-tuned models for direct application in downstream tasks. Often, the models finetuned by researchers are not made publicly available, limiting their utility outside the original research context. Additionally, even when such models are accessible, their usage frequently requires advanced programming skills, posing a significant barrier to their broader adoption in genomic research. Only a few CNN-based models come with associated web services, such as humanbase (https://humanbase.io/) which hosts Sei, DeepSEA, and Beluga to predict alterations of epigenetic features (10, 11), and the SpliceAI lookup (https://spliceailookup.broadinstitute.org) predicting splice site alterations SpliceAI (12). Their application, however, is confined to annotating .vcf files or short DNA sequences up to 1 kb. Moreover, they do not offer insights into the sequence determinants underlying the model’s predictions.

To bridge this gap, we introduce GENA-Web, a web service designed to offer a user-friendly platform for inferring sequence-based features using DNA transformer models. GENA-Web hosts models tailored for annotating promoters, splice sites, epigenetic features, and enhancer activities, as well as to highlight sequence determinants that underlying model prediction.

## MATERIAL AND METHODS

### Web service implementation

The front-end was implemented on TypeScript using React, Redux, Eslint, and Npm. To visualize genomic data, we integrated *igv*.*js* (13), ensuring an interactive and intuitive display of results. On the back-end, implemented using flusk and python, we structured each model as a separate Docker container, enabling isolation and scalability. These containers are designed to accept standardized inputs and provide outputs in genomic annotation formats, specifically .bed or .bedGraph. This modular architecture not only facilitates the maintenance and operation of the current system but also simplifies the integration of additional models in the future, enhancing the web server’s versatility and adaptability to evolving research needs.

### Pretraining of DNA Language Models

This study utilizes the pre-trained DNA language models DNABERT and GENA-LM, as introduced in (4) and (7), respectively. To ensure the manuscript’s completeness, we briefly describe the characteristics of these pre-trained models. DNABERT, which incorporates a vocabulary of DNA k-mers of length 6, underwent pretraining on the human hg38 genome assembly. The model is accessible through its official GitHub repository: https://github.com/jerryji1993/DNABERT.

The GENA-LMs collection includes models that vary in architecture, parameter count, and the datasets used for pre-training. Specifically, all GENA-LMs employed in our research were pre-trained using the human T2T genome assembly. For tokenization, GENA-LMs utilize the Byte Pair Encoding (BPE) strategy, resulting in tokens of varying lengths with a median of 8 bp.

Both the GENA-LMs and DNABERT models were subjected to masked language modeling (MLM) during pretraining. This process involved tokenizing segments of the human genome sequence, flanking them with CLS and SEP tokens. In alignment with the BERT pretraining protocol, 15% of the tokens in each segment were designated for prediction: 80% were masked (replaced by a special “MASK” token), 10% replaced with random tokens, and 10% left unchanged. The GENA-LMs underwent pretraining over 1-2 million steps with a batch size of 256.

Depending on the specific genomic task, we utilized one of the following GENA-LMs: *gena-lm-bert-base-t2t* (110M parameters, employing full attention, with an input length capacity of 4.5 kb), *gena-lm-bert-large-t2t* (336M parameters, also using full attention, with the same input length limit), or *gena-lm-bigbird-base-sparse-t2t* (110M parameters, utilizing sparse attention, and capable of processing inputs up to 36 kb).

### Fine-tuning process

Each task featured on the web server was developed through the process of fine-tuning, utilizing one of two publicly available pre-trained DNA language models: DNABERT (4) or GENA-LMs (7), as detailed in Supplementary Table 1. For the DNABERT model, we always use the *DNABERT6* model. The optimal GENA-LM for each specific task was selected based on performance metrics reported in (7). Supplementary Table 1 provides detailed information regarding the specific pre-trained model deployed for each downstream task, along with the performance scores achieved during the fine-tuning phase.

For splice sites annotation using the DNABERT model, we obtained very low performance scores (f1 score near 0.5). Therefore, this model was not included in the service. For promoter activity prediction, we used EPDnew-derived promoter sequences as positive samples set and random genomic sequences non-overlapping promoters as a negative samples set. For other tasks, we used original datasets provided by authors (10, 12, 14).

In the fine-tuning process of GENA-LM models, we start by tokenizing input sequences and adding CLS and SEP tokens at the beginning and end, respectively. Sequences are adjusted to fit model requirements through padding or truncation. These tokenized sequences feed into models built on the pre-trained GENA-LM architecture, which includes an additional fully-connected output layer sized according to the hidden units of the model and the specific target task.

For classification tasks, we use a softmax function for single-label tasks and a sigmoid function for multi-label tasks, along with appropriate loss functions: cross-entropy for the former and binary cross-entropy with logits for the latter. Regression tasks use mean squared error as the loss function without an activation function on the output layer. The hidden states of CLS tokens is used for sequence classification and regression, while all hidden states are utilized for token-level classification, like splice site prediction.

Both the weights of the final layer and the overall GENA-LM parameters were fine-tuned, with a learning rate warm-up applied. The ideal number of training and warm-up steps is empirically determined for each task.

In the fine-tuning process of the DNABERT model, minor modifications were made to the original source code to suit our tasks. The tokenization process utilized the model’s default tokenizer, which divides the sequence into tokens, each representing a k-mer of fixed length 6. While the default fine-tuning parameters were largely retained, adjustments were made to the learning rate and batch size to improve convergence. The dimensionality of the final fully-connected layer of the model was tailored to the demands of the given task, employing the same types of loss functions as those used with the GENA-LM models. This process involved the fine-tuning of both the weights of the final layer and the main body of the DNABERT model with no parameters being frozen.

### Long input processing

When the input sequence surpasses the model’s maximum input capacity, we divide the input into smaller, manageable chunks and independently compute predictions for each segment. Recognizing the significance of contextual information for precise predictions, our strategy involves employing overlapping chunks. This approach ensures that predictions are derived from the segment where the region of interest has the maximal contextual information. We employed this overlapping chunk methodology for tasks that, during training, were provided with extensive (*>*= 1000 bp) contextual information, such as chromatin annotation and splice site prediction. Conversely, for tasks where the contextual information during training was less extensive, we opted for non-overlapping chunks to increase computational efficiency. In all cases, collected predictions for all chunks are concatenated together into a single output file.

### Computation of Input Token Attribution Scores

Token attributions scores are calculated using the Layer Integrated Gradients method (15) implemented in the captum library (16).

## RESULTS

### Overview of the GENA-Web service

*Input Requirements* Users are prompted to provide the DNA sequence, select the desired model, and specify the task to be performed by the web service (Fig. 1, A). Presently, the service offers four distinct tasks for selection: promoter activity prediction, splice site annotation, prediction of epigenetic features, and enhancer activity assessment. Detailed descriptions of each task are provided in subsequent sections.

**Figure 1.**
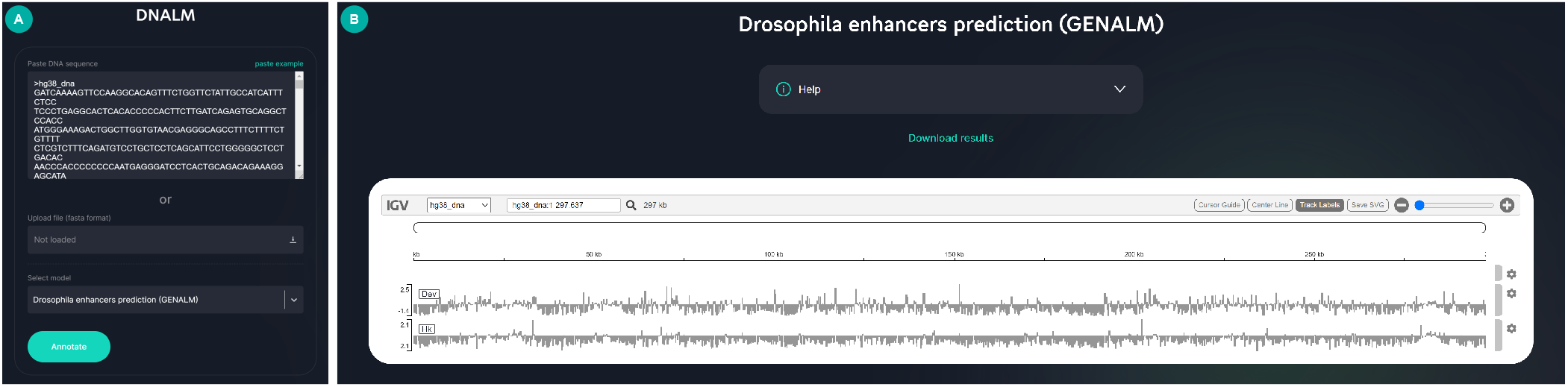
Input and output views of the GENA-Web service. A. Input screen. B. Example of service output page with IGV view and link to files download

The DNA sequence can be input directly into the designated field on the homepage or uploaded as a FASTA file. The input sequence length is currently limited to 1 megabase (Mb). Model and task selection are enabled by dropdown menus on the homepage.

*Output* The service generates DNA annotations as specified by the user, offering outputs both as downloadable files (using .*bed* format for qualitative features and .*bedGraph* for quantitative annotations) and through an interactive genome browser display (Fig. 1, B). Additionally, it provides insights into how specific elements of the input sequence influence the annotations. This analysis is conducted using the Integrated Gradients method (15), which assigns an “importance” score to each sequence token, highlighting its contribution to the feature annotation.

Users can view these token importance scores by clicking on a feature in the genome browser, which then displays the corresponding scores track. A high token importance score suggests that alterations to that token could significantly impact the annotated feature. The tokens have lengths of 6 bp for DNABERT and approximately 8 bp for GENA-LMs, allowing for a detailed examination of the sequence elements critical to the identified features.

## Tasks Available in GENA-Web Service

### GENA-Web currently offers four distinct tasks

#### Promoter Prediction

This task identifies potential promoter regions within the input sequence, providing a binary (yes/no) indication for segments of 300 bp in length. Given its training on human promoter sequences, its applicability might be limited to human or closely related mammalian genomes.

#### Splice Site Annotation

It involves classifying each sequence token based on its likelihood of representing a splice donor (SD) or acceptor (SA) site, with separate tracks for each. While trained on human data, we observed that this model demonstrates a capacity to identify splice site motifs across species, including distantly related organisms like *Drosophila*.

#### Epigenetic Profiling

This task offers predictions on DNA accessibility, histone modifications, and transcription factor bindings across various human cell lines, encompassing over 900 targers organized into several tracks for user convenience. Each reported target presents specific combination of cell type and epigenetic feature. For some cell types, predictions under treatment condition are also available. For example, the target named *K562* |*CTCF*| *None* reflects CTCF transcription factors binding in the untreated K562 human cells. More details about targets can be found in (10).

#### Enhancer Activity Annotation

Using outputs of the model trained on *Drosophila* cell reporter assays (14), this task evaluates the sequence’s potential to enhance the activity of housekeeping and developmental gene promoters, presented in two distinct tracks.

### Case Study I: Deciphering chimeric transcript structure with splice site prediction

In a recent study, we reported a complex structural variant (SV) on chromosome 9 (9p24) segregating across five generations within a family affected by congenital heart defects (Fig 2, A) (17). Detailed genomic analyses unraveled a chimeric gene formation, integrating the 5’ segment of the *KANK1* transcript with an intergenic segment located between *KANK1* and *DKK1*, and extending into the *DKK1* gene (Fig 2, A). Given the evidence of *KANK1* role in cardiomyocyte differentiation (18, 19), the emergence of this chimeric transcript, incorporating elements of *KANK1*, could potentially exhibit gain-of-function activity, contributing to the observed cardiac phenotype. However, the chimeric structure also encompasses an intergenic region, leaving it devoid of splice site annotations. This lack of information hindered the reconstruction of the mature mRNA sequence. Although *KANK1* is highly expressed in cardiac tissues, its minimal expression in blood made it challenging to directly study the transcript structure using RNA from the patient’s blood cells.

**Figure 2.**
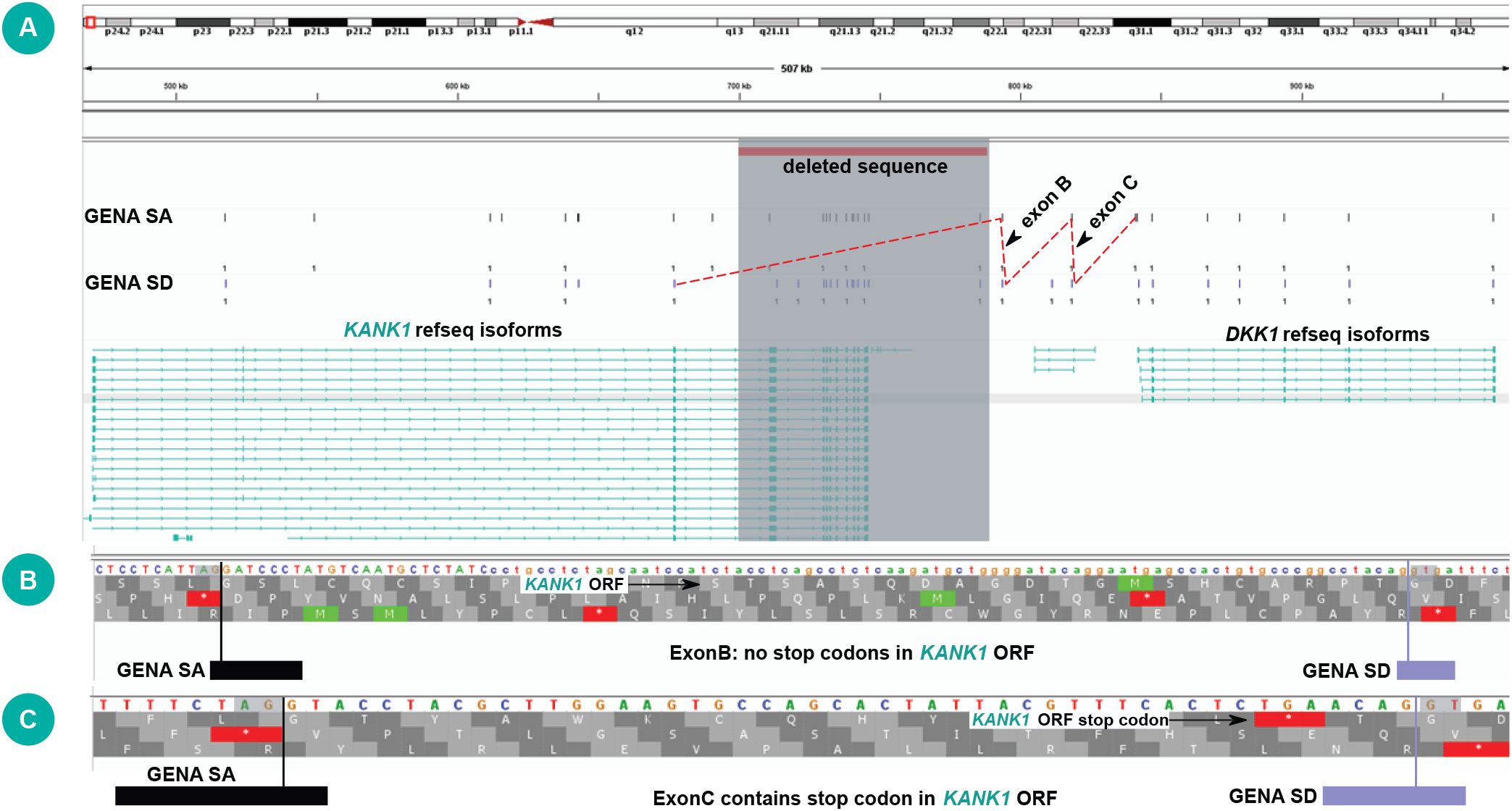
Inferring chimeric transcript structure using GENA-Web. A. The structure of *KANK1-DKK1* loci showing the deleted region and Splice Acceptor (SA) and Splice Donor (SD) sites predicted using GENA-Web (Splice Sites Annotation with GENA-LM task). B and C. Structure of exons predicted within the intergenic region. Exon boundaries are shown by vertical lines

To explore potential splice sites within the chimeric region, we employed GENA-Web-based splice sites annotation. This investigation identified two pairs of splice donor and acceptor sites within the intergenic segment, indicating the possible formation of two exons (illustrated in Fig.2, A for the whole transcript structure, and zoomed in Fig.2, B and C for the individual exons). The first exon (Fig.2, B) maintains the *KANK1* reading frame without introducing stop codons. However, the subsequent exon, spanning 48 nucleotides as shown in Fig. 2, C, introduces a stop codon within the *KANK1* reading frame. Consequently, this chimeric transcript likely contains a premature termination codon, subjecting it to nonsense-mediated RNA decay. Given that *KANK1* is not the haploinsufficient gene, these findings suggest that the *KANK1-DKK1* chimeric transcript does not play a direct role in the familial heart condition observed.

### Case Study II: Studying evolution of splice sites determinants

Further evaluating the capabilities of our GENA-LM web service, we tested the task of splice site annotation across different species. This examination focused on the service’s accuracy in identifying donor and acceptor sites within the genomes of human, mice, zebrafish, and fruit fly using the model fine-tunned exclusively on human data. Given this model s training background, its effectiveness in identifying splice sites in non-human species depends on the conservation of sequence determinants for splice donor and acceptor sites across these organisms.

To conduct this assessment, we sourced random gene sequences from the positive strand of each genome via the UCSC genome browser. These sequences were then processed through the GENA-LM web service to gather donor and acceptor sites. A prediction was considered accurate if the actual splice site was encompassed within the bounds of the predicted token’s range, emphasizing the model’s ability to pinpoint splice sites within a specified genomic interval. Our results (Fig. 3) demonstrated a high degree of accuracy in predicting donor and acceptor sites within human genes, achieving rates of 93% and 88%, respectively. The performance on the mouse and zebrafish genomes also yielded robust results, with donor site predictions achieving accuracies of 81% and 86%, and acceptor site predictions reaching 86% and 76%, respectively. However, the accuracy dropped when assessing the fruit fly genome, with only 54% accuracy for donor sites and 61% for acceptor sites. Nevertheless, Importantly, the accuracy rates observed for all species were markedly higher than those seen in a control scenario, wherein predictions from GENA-LMs were replaced with genomic sites selected at random. Such random selection resulted in an accuracy rate approaching 0%, with a p-value negligible when compared to the accuracy achieved by GENA-LM predictions. This toy example shows that there are elements of splice site grammar conserved among a range of animal taxa, aligning with findings previously documented in the literature (20).

**Figure 3.**
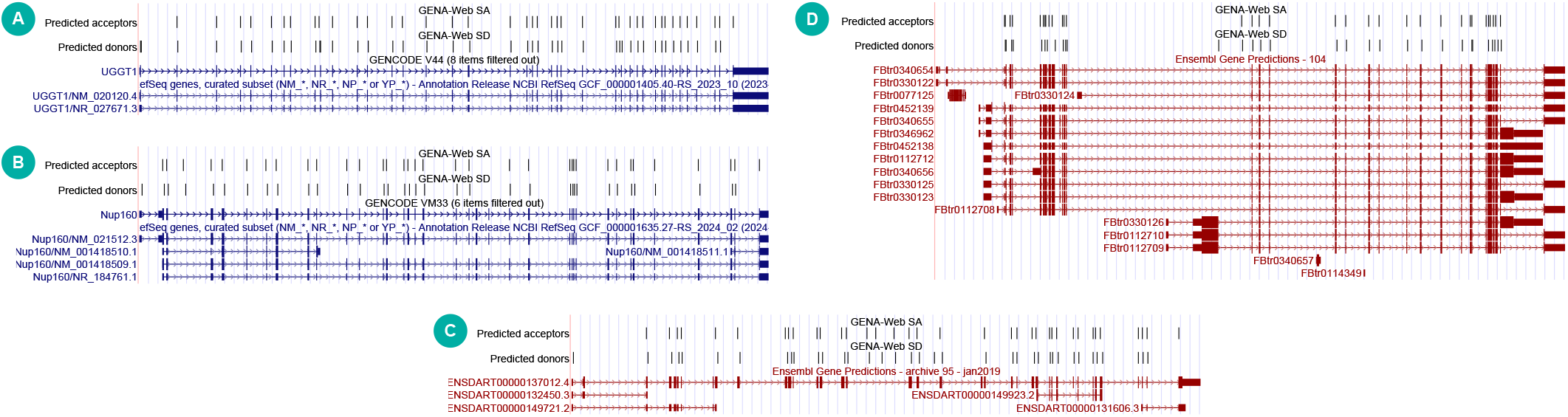
Cross-species inference of splice sites using a model trained on human data. Annotations of splice sites generated by GENA-Web are presented alongside screenshots from the UCSC Genome Browser. The species analyzed include A. *Homo sapiens* B. *Mus musculus* C. *Danio Rerio* D. *Drosophila Melanogaster*. SD - splice donor, SA - splice acceptor

## CONCLUSIONS AND FUTURE PERSPECTIVES

GENA-Web introduces a novel web service dedicated to annotating genomic data, capable of inferring approximately 1000 features directly from DNA sequences. Its capacity to process inputs up to 1 Mb in length enables comprehensive analyses and integrates seamlessly with models designed to leverage extensive contextual information (7). Despite the broad utility of GENA-Web, it’s important to note that a significant portion of its outputs pertains to chromatin states, primarily based on models trained with human data. Looking ahead, expanding GENA-Web’s functionality through the integration of new models and tasks will be a key area for development, enhancing its applicability across a wider range of genomic research areas.

## DATA AVAILABILITY

Source code is available on GitHub: https://github.com/AIRI-Institute/GENA_Web_service,https://github.com/AIRI-Institute/GENA_Web_docker. The models are available on HuggingFace (with references specified in Supplementary Table 1)

## AUTHOR CONTRIBUTIONS

A.S., M.P., and Y.K: models development. A.S. M.P. and D.P.: web-service implementation. N.A., M.T, S.M. - front-end development. V.F., M.B., O.K.: Conceptualization, Formal analysis, Methodology, Validation. V.F.: Writing—original draft. All authors: Writing—review & editing.

## ACKNOWLEDGEMENTS

The case study was supported by RSF grant 22-14-00247. Veniamin Fishman acknowledges the support of the Institute of Cytology and Genetics SB RAS core facilities for providing computational resources (project 121031800061-7, Mechanisms of genetic control of development, physiological processes and behavior in animals)

