## supplementary table 1 for "GENA-Web - GENomic Annotations Web Inference using DNA language models"

Supplementary table 1: Tasks implemented in the web-service. For chromatin annotation we report three scores, each overaged accross features in all accessed cell types: predictions of transcription factor binding sites (TF), DNAse I hypersensitivity sites (DS), and histone modifications (HM). For *Drosophila* enhancer activity prediction we provide separate scores for houskeeping (hs) and developmental (dev) enhancer activity

| Downstream task | Task type (number of targets) | Metric | DNABERT | GENA-LM | Loss type | Model availability | |
|---|---|---|---|---|---|---|---|
| | | | | | | GENA-LM source | DNABERT source |
| Promoter activity prediction | classification (2 classes) | F1 | 83.36 | **85.26** | Cross-entropy | https://huggingface.co/AIRI-Institute/gena-lm-bert-large-t2t/tree/gena_web_promoters_2000 | https://huggingface.co/AIRI-Institute/DNABERT/tree/gena_web_dnabert_promoters_2000 |
| Splice sites classification | classification (3 classes) | AUC | — | **94.78** | Binary Cross-entropy | https://huggingface.co/AIRI-Institute/gena-lm-bigbird-base-t2t/tree/gena_web_spliceai | |
| Chromatin annotation (TF) | classification (565 classes) | | 96.37 | **96.54** | | | |
| Chromatin annotation (DHS) | classification (125 classes) | AUC | 92.7 | **92.83** | Binary Cross-entropy | https://huggingface.co/AIRI-Institute/gena-lm-bert-base-t2t/tree/gena_web_deepsea | https://huggingface.co/AIRI-Institute/DNABERT/tree/gena_web_dnabert_deepsea |
| Chromatin annotation (HM) | classification (104 classes) | | 86.13 | **86.64** | | | |
| Drosophila enhancer prediction (dev) | regression (1 target) | PCC | **70.2** | 65.1 | MSE | https://huggingface.co/AIRI-Institute/gena-lm-bert-base-t2t/tree/gena_web_deepstarr | https://huggingface.co/AIRI-Institute/DNABERT/tree/gena_web_dnabert_deepstarr |
| Drosophila enhancer prediction (hs) | regression (1 target) | | **77.9** | 75.9 | | | |
